# Memory quality modulates the effect of aging on memory consolidation during sleep: Reduced maintenance but intact gain

**DOI:** 10.1101/547448

**Authors:** Beate E. Muehlroth, Myriam C. Sander, Yana Fandakova, Thomas H. Grandy, Björn Rasch, Yee Lee Shing, Markus Werkle-Bergner

**Author notes:** Correspondence concerning this article should be addressed to: Beate E. Muehlroth or Markus Werkle-Bergner; Max Planck Institute for Human Development, Lentzeallee 94, 14195 Berlin, Germany.

## Abstract

Successful consolidation of associative memories relies on the coordinated interplay of slow oscillations and sleep spindles during non-rapid eye movement (NREM) sleep, enabling the transfer of labile information from the hippocampus to permanent memory stores in the neocortex. During senescence, the decline of the structural and functional integrity of the hippocampus and neocortical regions is paralleled by changes of the physiological events that stabilize and enhance associative memories during NREM sleep. However, the currently available evidence is inconclusive if and under which circumstances aging impacts memory consolidation. By tracing the encoding quality of single memories in individual participants, we demonstrate that previous learning determines the extent of age-related impairments in memory consolidation. Specifically, the detrimental effects of aging on memory maintenance were greatest for mnemonic contents of medium encoding quality, whereas memory gain of weakly encoded memories did not differ by age. Using multivariate techniques, we identified profiles of alterations in sleep physiology and brain structure characteristic for increasing age. Importantly, while both ‘aged’ sleep and ‘aged’ brain structure profiles were associated with reduced memory maintenance, inter-individual differences in neither sleep nor structural brain integrity qualified as the driving force behind age differences in sleep-dependent consolidation in the present study.

## Introduction

Non-rapid eye movement (NREM) sleep is critical for the long-term retention of memories (Rasch and Born 2013). Most studies, so far, have focused on younger adults and provided compelling evidence that memory consolidation during sleep is supported by the interplay of slow oscillations and sleep spindles that guide the transformation of labile hippocampus-dependent memories into durable neocortical representations (Born and Wilhelm 2012; Diekelmann and Born 2010; Gais and Born 2004; Walker 2009). Corresponding research in older adults, though, has produced inconsistent findings (Scullin 2013; Sonni and Spencer 2015; Wilson et al. 2012; for a review see Scullin and Bliwise 2015). To date, the available evidence is inconclusive as to whether and under which circumstances memory consolidation is affected by advancing age.

This inconsistency might, in parts, arise from unknown variations in the *quality* of individual memories. The depth of encoding (Craik and Lockhart 1972) and degree of learning (Tulving 1967) discriminate memories with regard to their *memory strength* or *memory quality*. In general, insufficient and shallow processing of new information appears to hamper learning success in old age (Craik et al. 2010; Naveh-Benjamin et al. 2007; Shing et al. 2008). Further, as older adults have deficits in binding items into cohesive and distinct memory representations (Naveh-Benjamin 2000; Old and Naveh-Benjamin 2008; Shing et al. 2008; St-Laurent et al. 2014), the quality of newly formed memories might be critically reduced. Crucially, it is likely that sleep-dependent memory consolidation does not affect all encoded memories in the same way (Diekelmann et al. 2009; Schoch et al. 2017; Stickgold and Walker 2013). Several studies imply that memories of intermediate quality are prioritized during sleep, resulting in greater benefits for weakly encoded memories (Diekelmann et al. 2009; Drosopoulos et al. 2007; Kuriyama et al. 2004; Schapiro et al. 2018; Stickgold 2010; but see Tucker and Fishbein 2008). Hence, proper assessment of age differences in sleep-dependent memory consolidation requires consideration of the quality of encoded memories.

Already 50 years ago, Tulving called for an improvement of behavioral memory measures (Tulving 1964, 1967) arguing that *“the ‘average’ item is a highly abstract and elusive entity having no readily identifiable counterparts in the empirical realm”* (Tulving 1967, p. 183). Still, research has only recently started to address this problem with regard to memory consolidation. By looking at each memory’s individual “fate” it becomes possible to identify whether and how the success of memory encoding influences later consolidation processes (Dumay 2016; Fenn and Hambrick 2013): sleep potentially stabilizes previously successfully encoded memories but may also enhance the availability of initially poor memories above a pre-sleep learning level (Ellenbogen et al. 2006; Nettersheim et al. 2015). So far, prevailing evidence speaks for a primary role of sleep in *memory maintenance*: by passively protecting memories against interference (Wixted, 2004) and actively reactivating and stabilizing memory engrams (Rasch et al. 2007), memories are maintained across sleep (Dumay 2018; Fenn and Hambrick 2013; Schreiner et al. 2018). Behaviorally observed *memory gains* that reflect the availability of initially poor memories during later memory retrieval appear to rely less on sleep (Dumay 2018; Fenn and Hambrick 2013; Schreiner et al. 2018).

Successful consolidation during sleep not only requires successful memory encoding. Brain atrophy in old age directly affects memory-relevant regions in the medial temporal lobe (MTL) (Raz et al. 2005), potentially leading to impaired encoding, retrieval, and importantly, to deficient consolidation of memories (Craik and Rose 2012; Shing et al. 2011, 2008; Ward et al. 1999; Werkle-Bergner et al. 2006). Moreover, age-related structural changes in brain regions involved in slow oscillation and spindle generation have frequently been related to age-related sleep alterations (Dubé et al. 2015; Fogel et al. 2017; Landolt and Borbély 2001; Mander et al. 2013; Varga et al. 2016). Accordingly, age-related brain atrophy might lead to worse memory consolidation in old age, by impairing the neurophysiological underpinnings of successful consolidation (Helfrich et al. 2018; Mander et al. 2013; Muehlroth et al. in press).

Indeed, aging involves substantial changes in sleep physiology that affect NREM sleep in particular (Mander et al. 2017). Compared to younger adults, in old age the deepest NREM sleep stage, so-called slow-wave sleep (SWS), is strikingly reduced (Carrier et al. 2011; Mander et al. 2017; Ohayon et al. 2004) and slow oscillations and sleep spindles themselves are less often observed (Crowley et al. 2002; Dubé et al. 2015; Fogel et al. 2012). These age-related changes in NREM sleep may affect memory consolidation and account for dysfunctional protection of memories against forgetting in old age (Baran et al. 2016; Buckley and Schatzberg 2005; Cherdieu et al. 2014; Harand et al. 2012; Hornung et al. 2005; Varga et al. 2016; but see Aly and Moscovitch 2010; Backhaus et al. 2007; Wilson et al. 2012).

In the present study we tracked the learning history of single scene–word associations within individual younger and older adults. This procedure allowed us to test how encoding quality modulates age differences in memory consolidation during sleep. Using a multivariate statistical approach, we then examined how a senescent sleep profile, that is marked by reductions in both slow oscillations and sleep spindles, relates to age differences in overnight memory consolidation. Finally, using the same multivariate approach, we addressed how structural decline in specific consolidation- and memory-relevant brain regions contributes to alterations in both sleep and overnight changes in memory performance.

## Materials and methods

### Participants and procedure

#### Participants

Overall, 34 healthy younger adults (19–28 years) and 41 healthy older adults (63–74 years) took part in the experiment. Data from one older adult were excluded due to an incidental finding during magnet resonance imaging (MRI). Four younger and four older adults had to be excluded due to technical failures during data collection, resulting in a final sample of 30 younger (*M*_age_ = 23.7 years, *SD*_age_ = 2.6; 17 females) and 36 older adults (*M*_age_ = 68.92 years, *SD*_age_ = 3.04; 16 females) for the behavioral analyses. Since parts of some participants’ neural data (polysomnography [PSG] or MRI) were missing or of bad quality, final PSG analyses were conducted with 24 younger (*M*_age_ = 23.61 years, *SD*_age_ = 2.55; 13 females) and 31 older adults (*M*_age_ = 68.63 years, *SD*_age_= 3.10; 15 females). From this sample, two older adults had to be excluded for structural MRI analyses resulting in a sample of 29 older adults for the respective analyses (*M*_age_ = 68.64 years, *SD*_age_ = 3.10; 14 females).

All participants were right-handed native German speakers with no reported history of psychiatric or neurological disease, or any use of psychoactive medication. All older adults completed the Mini-Mental State Examination (MMSE; *M* = 29.24, *SD* = 1.12, range: 26–30; Folstein et al. 1975) and passed a brief memory screening before inclusion in the final experiment. General subjective sleep quality was controlled by assessing the Pittsburgh Sleep Quality Index (PSQI; Buysse et al. 1989). The study was approved by the Ethics Committee of the *Deutsche Gesellschaft für Psychologie* (DGPs) and conducted at the *Max Planck Institute for Human Development* in Berlin. All participants gave written consent to their participation in the experiment after being informed about the complete study procedure.

#### General procedure

At the core of the experimental design was an associative memory paradigm that consisted of a learning session on the first day (Day 1) as well as a delayed cued recall task approximately 24 hours later (Day 2) (see Figure 1 for illustration of the study procedure). During the nights before and after learning (experimental nights PRE and POST), sleep was monitored at the participants’ homes using ambulatory PSG. Prior to the first experimental night an adaptation night familiarized the participants with the PSG procedure. Structural MRI data were collected on Day 2. Furthermore, electroencephalography (EEG) was recorded during learning on Day 1. Additionally, functional MRI data were collected during delayed recall on Day 2. Neither the EEG nor the fMRI data are included in the present report (but see Sommer et al. 2019 and Sander et al. 2019 on oscillatory mechanisms and neural pattern similarity during memory encoding that allow for successful memory formation).

**Figure 1.**
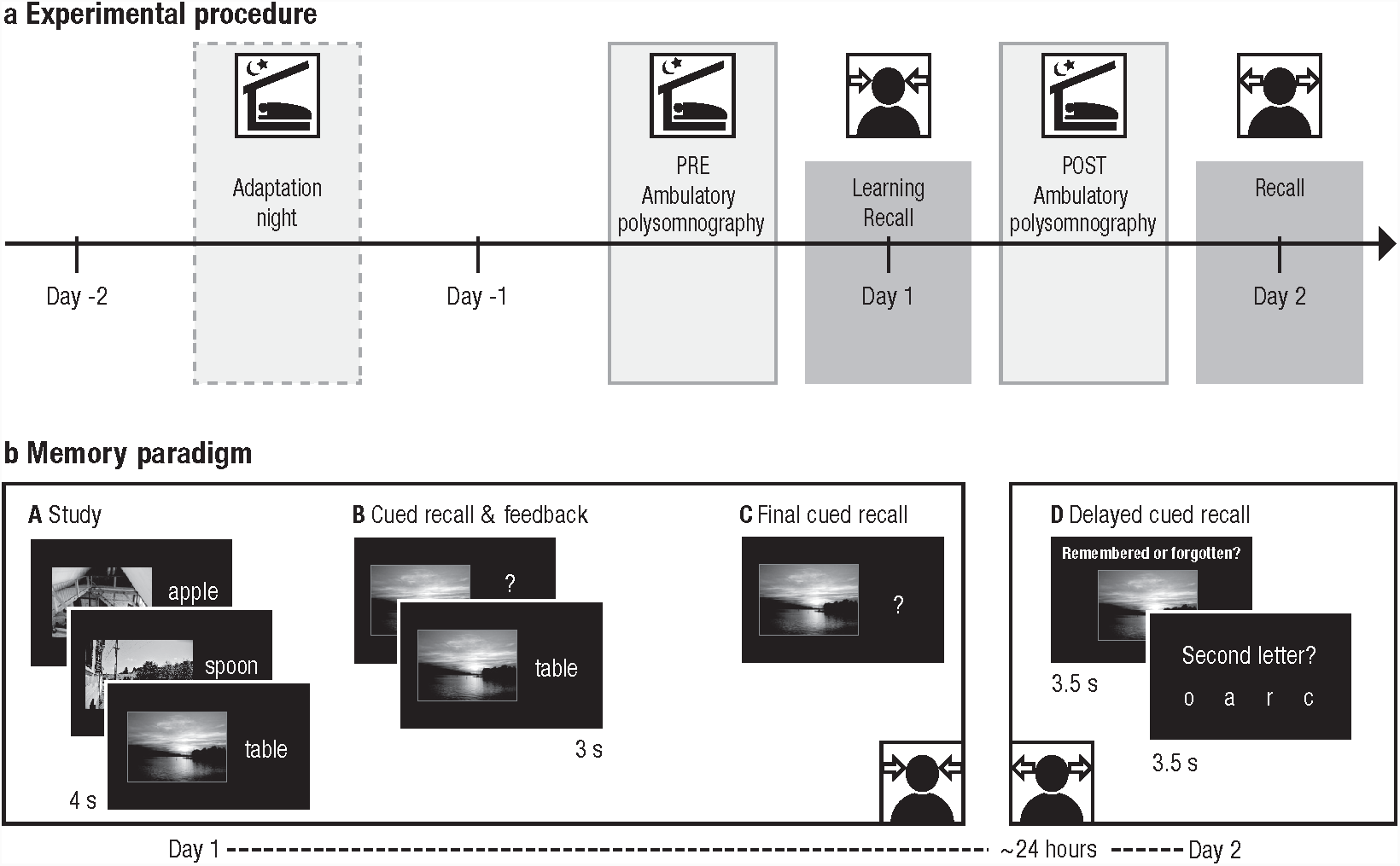
a. Experimental procedure (reproduced from Muehlroth et al. in press). The memory task at the core of the experiment consisted of a learning phase and an immediate recall on Day 1 as well as a delayed recall approximately 24 hours later. Sleep was monitored in the nights before (PRE) and after (POST) learning using ambulatory polysomnography. A prior adaptation night familiarized the participants with the sleep recordings. b. Memory paradigm. (A) During study, participants were instructed to remember 440 (younger adults) or 280 scene–word pairs (older adults). (B) During the cued recall and feedback phase the scene was presented as a cue to recall the corresponding word. Irrespective of recall accuracy, the original pair was presented again to allow for re-study. The whole cued recall and feedback cycle was performed once in younger and twice in older adults. (C) During final recall, scenes again served as cues to recall the corresponding word, but no feedback was provided. (D) Delayed cued recall took place approximately 24 hours later. Participants were presented with the scenes only and had to indicate if they still remembered the associated word. Afterwards they had to select the corresponding second letter of the word to verify their true memory of the associate.

#### Memory paradigm

The memory task and the stimulus set are described by Fandakova et al. (2018) and Muehlroth et al. (in press) in detail. During initial study, randomized combinations of scenes and concrete nouns were presented on a black screen for 4000 ms. Participants were instructed to remember these scene–word pairs using a previously trained imagery learning strategy. During the ensuing cued recall phase, scenes served as cues for participants to verbally recall the associated word. Independent of recall accuracy, the correct scene–word pair was presented again for 3 seconds and participants were encouraged to restudy the combinations. At the end of Day 1, participants completed a final cued recall test without feedback. Task difficulty of the encoding task was adjusted between the age groups to achieve comparable recall success: First, younger adults learned 440 pairs, whereas older adults learned 280 pairs on Day 1. Second, younger adults completed one cued recall block with feedback, whereas older adults completed an additional cued recall block with feedback that was excluded from further analyses (*Md*_performance_ = 3.93 %).

Delayed cued recall of the scene–word pairs took place approximately 24 hours later. Both younger and older adults were presented with 280 scenes for 3500 ms. Via keypress, participants had to indicate whether they still remembered the respective word (“remembered” vs. “forgotten”). Afterwards they had to select the corresponding second letter of the word out of four letter options to verify their true memory of the associate. Importantly, for the older age group, all of the 280 studied pairs were presented. For younger adults, items were chosen with regard to their learning history. If permitted by the individual learning performance on Day 1, this resulted in a selection of pairs, half of which had been recalled in the criterion cued recall the day before (Supplementary Figure 1 for inter-individual differences in trial composition during delayed recall on Day 2). All behavioral measures were adjusted for inter-individual differences in learning trajectories on Day 1 (see below).

#### Behavioral analyses

For each recall phase during the learning task on Day 1 the number of correctly recalled items was calculated and divided by the corresponding number of trials. Answers during delayed recall on Day 2 were only classified as being correct if participants both responded to remember the word *and* if they indicated the true letter afterwards. Reaction times for giving the “remember vs. forgotten” judgement were extracted for all correct trials.

Given the nature of the memory task with several recall phases, we were able to analyze recall success on Day 2 as conditional on the recall success on Day 1 (Figure 2a, cf. Dumay 2016). We thus distinguished two categories of items: (a) *maintained items* (items recalled both during final cued recall on Day 1 and during delayed recall on Day 2), (b) *gained items* (items recalled during delayed recall on Day 2, but not during final cued recall on Day 1). Maintained items were further divided into those items that were recalled during both recall phases on Day 1 (= *high memory quality*) and into those that were only successfully recalled during final cued recall (= *medium memory quality*). Gained items that were not at all recalled on Day 1 were considered having a *low memory quality* with respect to encoding on Day 1 (Figure 2a). Within each of the resulting three categories we determined the probability of recalling an item during delayed recall on Day 2. All analyses focused on successful memory recall on Day 2. Hence, the complementary categories of items not recalled on Day 2 were not included in further analyses.

**Figure 2.**
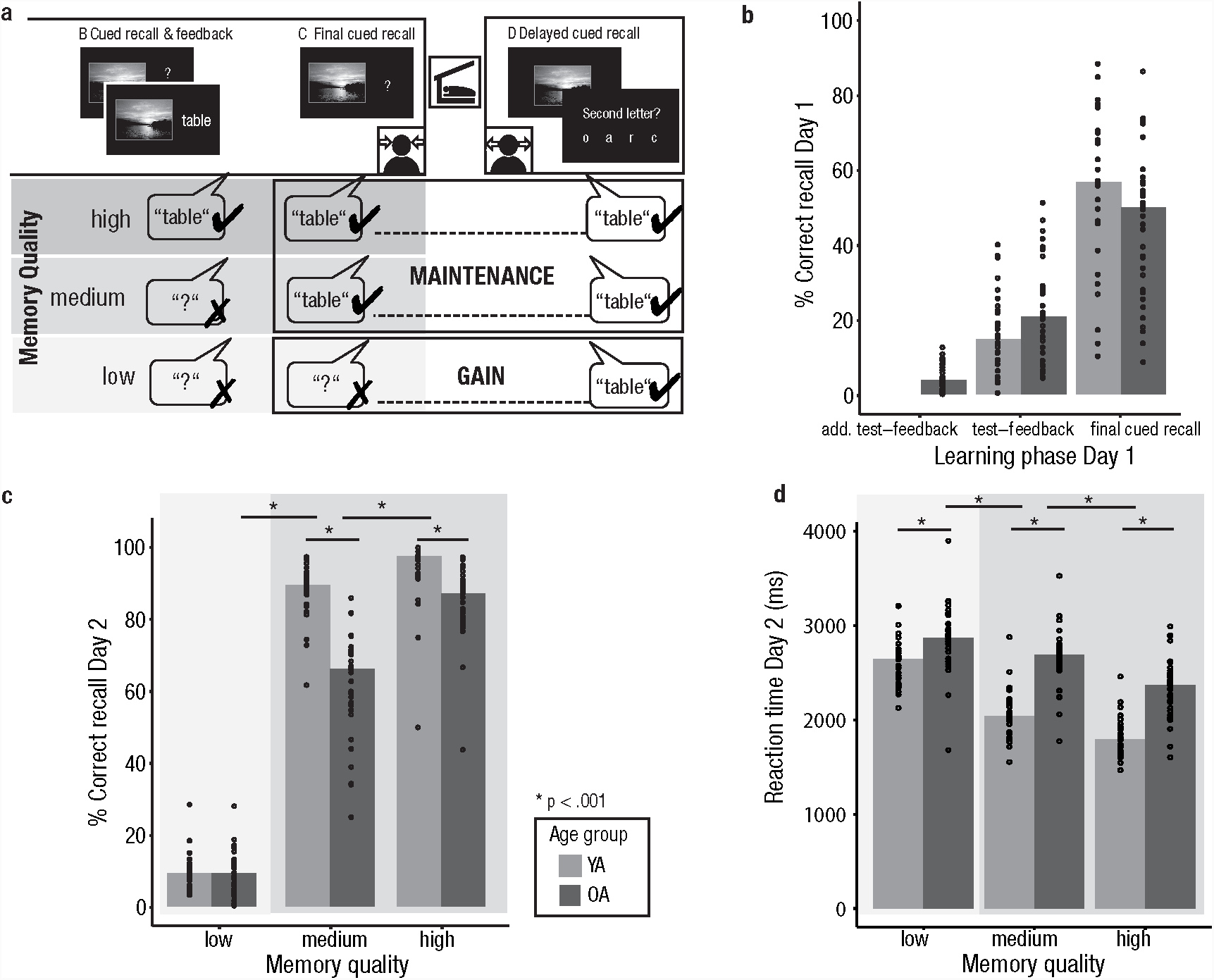
Memory quality limits the effects of aging on memory consolidation. (a) Identification of memory quality. The recall success of items during delayed recall on Day 2 was analyzed based on learning success on Day 1. Recall success during final cued recall on Day 1 determined whether an item was *gained* (shaded in light gray) or *maintained* across sleep (shaded in darker gray). Recall success of the test–feedback phase conditioned memory quality. (b) Learning performance on Day 1. The learning performance of both younger (light gray) and older adults (dark gray) increases across repeated learning cycles and is comparable in both age groups. Bars indicate group medians, dots mark the performance of individual participants. In older adults, an additional test-feedback cycle with low initial recall success was performed. (c) Recall success on Day 2 increases with memory quality (from left to right on the x-axis). Younger and older adults show a similar memory gain across sleep (shaded in light gray), but memory maintenance is reduced in older compared to younger adults (shaded in darker gray). The maintenance of medium-quality items is particularly impaired. (d) Similar to memory performance, reaction times during delayed recall on Day 2 are conditioned on memory quality. YA: younger adults; OA: older adults.

Behavioral analyses were performed using non-parametric statistical tests, as not all memory measures met the assumptions for normality. Due to its robustness against violations of the normal distribution (Glass et al. 1972) and the lack of an adequate non-parametric equivalent, mixed factorial ANOVAs with the between factor AGE GROUP and the within factor MEMORY QUALITY were calculated for both recall success and reaction times on Day 2 to test for differences in memory retrieval. Post-hoc testing was conducted calculating non-parametric Mann-Whitney *U* Tests for independent samples and Wilcoxon signed-rank tests for matched pairs.

### Sleep EEG data acquisition and analyses

#### Data acquisition

During the two experimental nights before (PRE) and after learning (POST), sleep was recorded using an ambulatory PSG device (SOMNOscreen plus; SOMNOmedics, Germany). Eight scalp electrodes were attached according to the international 10–20 system for electrode positioning (Jasper 1958; Oostenveld and Praamstra 2001) (Fz, F3, F4, C3, C4, Cz, Pz, Oz) along with two electrodes on the mastoids A1 and A2 that later served as the offline reference. All impedances were kept below 6 kO. Data were recorded using Cz as the online reference for all EEG derivations and AFz as ground. EEG channels were recorded between 0.2–75 Hz with a sampling rate of 128 Hz. Additionally, a bilateral electrooculogram (EOG) was assessed. Two submental electromyogram channels (EMG) were positioned left and right inferior of the labial angle and referenced against one chin electrode. Electrical activity of the heart was recorded using two electrocardiogram channels (ECG).

#### EEG pre-processing

Data was preprocessed using *BrainVision Analyzer 2*.*1* (Brain Products, Germany), *Matlab R2014b* (Mathworks Inc., Sherborn, MA), and the open-source toolbox *Fieldtrip* Oostenveld et al. *(2011)*. *All EEG channels were re-referenced against the average of A1 and A2*. *Afterwards*, *sleep was visually scored on 30-second epochs according to the standard criteria suggested by the American Academy for Sleep Medicine* (AASM; Iber et al. 2007). Strong body movements were marked and bad EEG channels were visually rejected. For the remaining channels and time points, automatic artefact detection was implemented on 1-second long segments (see Muehlroth et al. in press, for more details). Global sleep parameters were estimated based on the visually scored sleep data. Total sleep time (TST) was calculated as time spent in stage 1, 2, SWS, and rapid eye movement (REM) sleep. Wake after sleep onset (WASO) was defined as the proportion of time awake between sleep onset and final morning awakening. Sleep latency described the timespan from lights off, as documented by participants, to the first occurrence of stage 2, SWS, or REM sleep in minutes. Finally, sleep efficiency was estimated by dividing TST by the time between sleep onset and final morning awakening.

#### Power spectral analysis

To get more fine-grained indicators of neural processes during deep NREM sleep, power spectral analyses of the sleep EEG data were conducted. Fast Fourier Transformation (FFT) with frequency limits between 0.5 and 30 Hz was applied on 5-second intervals using a Hanning function with no overlap. Slow-wave activity (SWA), defined as spectral power in the delta frequency range (0.5–4.5 Hz), was calculated for all NREM epochs (including stage 2 and SWS) and, to minimize confounding effects like skull thickness, normalized to the total power of the whole spectrum. As in previous studies frontal SWA was found to be particularly significant for cognitive functions of sleep (e.g., Mander et al. 2013), we focused on frontal SWA as the average between the two frontal derivations F3 and F4.

#### Spindle and slow oscillation detection

Slow oscillations and spindles were detected during NREM epochs (i.e., stage 2, and SWS) using established algorithms (spindles: Klinzing et al. 2016, 2018; Mölle et al. 2011; slow oscillations: Mölle et al. 2002; Ngo et al. 2013). The algorithms are described in detail in Muehlroth et al. (in press). In short, spindle detection was based on the smoothed root-mean-square (RMS) representation of the band-pass filtered EEG times series (6th-order Butterworth filter; slow spindles: 9–12.5 Hz; fast spindles: 12.5–16 Hz; cf. Cox et al. 2017). To account for individual differences in EEG amplitude, we anchored spindle identification on individually determined amplitude thresholds (Coppieters’t Wallant et al. 2016). Spindles were tagged if the amplitude of the smoothed RMS signal exceeded its mean by 1.5 SD of the filtered signal for 0.5 to 3 seconds. Spindles were merged if their boundaries were closer than 0.25 seconds and if the resulting spindle event remained within the time limit of 3 seconds.

For the detection of slow oscillations, the signal was band-pass filtered between 0.2 and 4 Hz at frontal electrodes (6th-order Butterworth filter). Putative slow oscillations were succeeding negative and positive half waves, separated by a zero-crossing, with a frequency between 0.5 and 1 Hz. Adaptive amplitude thresholds were defined separately for each participant: Slow oscillations had to exceed a trough potential of 1.25 times the mean trough potential of all putative slow oscillations and an amplitude of 1.25 times the average amplitude of all potential slow oscillations.

In line with the previously reported topography of slow oscillations, slow and fast spindles (Klinzing et al. 2016; Mander et al. 2017), we focused our analyses on frontal slow oscillations and slow spindles as well as central fast spindles. Only artefact-free slow oscillations and spindles were considered in the reported analyses. The density of slow oscillations and spindles was estimated by dividing the number of detected artefact-free events by the analyzed NREM sleep time (in minutes).

#### Statistical analyses

Sleep was assessed on two occasions (*before* and *after* learning). To investigate the variability of sleep architecture and physiology across age groups and experimental nights, we used mixed factorial ANOVAs with the within factor TIME and the between factor AGE GROUP. As not all sleep variables used in our analysis followed a normal distribution, post-hoc testing was conducted by calculating non-parametric Mann-Whitney *U* Tests for independent samples and Wilcoxon signed-rank tests for matched pairs. Median and quartile values of the variables were reported. In general, significance levels were set to α =.05 and tested two-sided. Statistical significance was controlled for multiple comparisons by Bonferroni-correcting the α-value and dividing it by the number of performed comparisons.

### MRI data acquisition and structural MRI analyses

Whole-brain MRI data were acquired with a Siemens Magnetom 3T TimTrio machine (high-resolution T1-weighted MPRAGE sequence: TR = 2500 ms, TE = 4.77 ms, FOV = 256 mm, voxel size = 1×1×1 mm^3^). Estimates of brain volume in regions of interest (ROI) were derived by means of voxel-based morphometry (VBM) using statistical parametric mapping software (SPM12, http://www.fil.ion.ucl.ac.uk/spm), the Computational Anatomy Toolbox (CAT 12, http://www.neuro.uni-jena.de/cat), and the REX toolbox (http://web.mit.edu/swg/rex/rex.pdf) (see Muehlroth et al. in press for a detailed description of the analyses steps). All measures were adjusted for differences in total intracranial volume (cf. Raz et al. 2005). Bilateral medial prefrontal cortex (mPFC), thalamus, entorhinal cortex, and hippocampus were selected as ROIs due to their involvement in memory consolidation and in the generation of spindles and slow oscillations (Gais and Born 2004; Nir et al. 2011; Steriade 2006). The mPFC mask was kindly provided by Bryce A. Mander (cf. Mander et al. 2013). All other ROIs were defined using the WFU PickAtlas toolbox (http://fmri.wfubmc.edu/software/pickatlas).

### Using Partial Least Squares Correlation (PLSC) to extract an age-specific latent sleep and brain structure profile

Slow oscillations and sleep spindles are not discrete entities but rather interact to enable memory consolidation (Helfrich et al. 2018; Latchoumane et al. 2017; Maingret et al. 2016). Aging results in global sleep alterations, affecting slow oscillations and spindles among other neurophysiological indicators (Carrier et al. 2011; Mander et al. 2017; Ohayon et al. 2004). To account for the interdependency of sleep processes and their joint age-related alterations, we applied *Partial Least Squares Correlation* (PLSC; Haenlein and Kaplan 2004; Krishnan et al. 2011) to extract a latent variable (LV) capturing age-related differences in various neurophysiological indicators of memory consolidation during sleep (namely frontal SWA, frontal slow oscillation density, frontal slow spindle density, and central fast spindle density).

First, a correlation matrix between the *Z*-standardized sleep variables (an n × 4 matrix) and chronological AGE (an n × 1 vector) was computed across all participants. Using singular value decompositions (SVD), the resulting correlation matrix was then decomposed into three matrices (UΔV^T^) based on which one LV was extracted in a least-squares sense. The resulting LV reflects the specific pattern of inter-individual differences in sleep measures that shares the largest amount of variance with inter-individual differences in the participants’ age. Statistical significance of the extracted LV was determined by 5000 permutations tests of its singular values. The reliability of the calculated LV weights (V) of each sleep parameter was tested using 5000 bootstrap samples. Dividing the weights by the bootstrap standard error provides bootstrap ratios (BSR) comparable to *Z*-scores. Hence, we considered an absolute BSR of 1.96 or higher as reliable, which approximately corresponds to a 95 % confidence interval. Finally, by projecting the original matrix of sleep parameters back onto the respective LV weights, we extracted a *latent sleep profile score* (LSPS) for each participant that reflects the degree to which each participant expressed the estimated robust latent sleep profile.

A comparable procedure was chosen to define a LV of brain structure, reflecting age-related gray matter decline in brain regions relevant for both sleep and memory (Gais and Born 2004; Nir et al. 2011; Steriade 2006). We computed a PLSC between an n × 4 matrix containing ROI volumes (i.e., mPFC, thalamus, entorhinal cortex, and hippocampus) and an n × 1 vector containing AGE to extract *latent brain structure score* (LBSS) capturing each participant’s manifestation of age-dependent loss in grey-matter volume across the four ROIs.

To relate the latent sleep profile and brain structure scores to the behavioral measures of memory gain and maintenance, Spearman’s rank-order correlation coefficients were calculated. Correlation coefficients were computed across the whole sample as well as separately within each age group. Fisher’s *Z*-transformation was used to test the null hypothesis of equal correlation coefficients in both the younger and the older sample.

All statistical analyses reported in this paper were conducted using *Matlab R2014b* (Mathworks Inc., Sherborn, MA), the open-source toolbox *Fieldtrip* (Oostenveld et al. 2011), *and RStudio 1*.*0*.*53* (RStudio, Inc., Boston, MA). The custom code and data necessary to reproduce all statistical results, figures, and supplementary material of this article are available on the *Open Science Framework repository* (https://osf.io/w76f3/).

## Results

### The effect of aging on memory consolidation is modulated by memory quality

In line with our attempt to adjust the encoding difficulty of the memory task between age groups (cf. Materials and methods), the percentage of correctly recalled scene–word associations did not differ systematically between age groups in the final cued recall on Day 1 and the preceding test-feedback phase (test feedback: *Z* = –1.67, *p* =.096; *Md*_YA_ = 14.77 %; *Md*_OA_ = 20.89 %; final cued recall: *Z* = 1.71, *p* =.087; *Md*_YA_ = 56.82 %; *Md*_OA_ = 50.00 %; Figure 2b). However, each item’s learning success on Day 1 (cf. Dumay 2016; Figure 2a) and the participants’ age systematically modulated recall success of items during delayed recall on Day 2 (Figure 2c). A mixed measures ANOVA with the between factor AGE GROUP and the within factor MEMORY QUALITY yielded significant main effects for both factors (MEMORY QUALITY: *F*(2, 128) = 2224.12, *p* <.001; AGE GROUP: *F*(1, 64) = 41.38, *p* <.001). Overall, successful recall was more likely when memory quality was high and, in general, younger adults had a higher probability of success during delayed recall. Post-hoc tests revealed that both age groups showed an equal rate of gained low-quality memories between Day 1 and Day 2 (*Z* = –0.08, *p* =.933; *Md*_YA_ = 9.12 %; *Md*_OA_ = 9.16 %). In contrast, older adults maintained a lower percentage of medium and high-quality memories than younger adults did (medium quality: Z = 6.40, *p* <.001; *Md*_YA_ = 89.64 %; *Md*_OA_ = 66.35 %; high quality: *Z* = 5.15, *p* <.001; *Md*_YA_ = 97.69 %; *Md*_OA_ = 87.28 %). The AGE × MEMORY QUALITY interaction was statistically relevant (*F*(2, 128) = 50.11, *p* <.001). Whereas both younger and older adults recalled less medium-quality than high-quality memories (*Z* = 6.54, *p* <.001), in comparison, older adults showed greater overnight loss for medium-quality memories (*Z* = –9.75, *p* <.001).

The effect of memory quality on delayed recall was also reflected in the corresponding reaction times during the delayed recall task on Day 2 (Figure 2d). Overall, older adults were slower in indicating that an item was remembered (*F*(1,64) = 49.03, *p* <.001) and in both age groups reaction times improved with higher memory quality (*F*(2,128) = 315.26, *p* <.001). Again, we observed a significant AGE × MEMORY QUALITY interaction (*F*(2,128) = 22.61, *p* <.001) with smaller, though significant (*Z* = –3.67, *p* <.001), age differences in reaction times for memories of low quality compared to both medium- and high-quality pairs. In line with the pronounced loss of medium-quality memories in old age, compared to younger adults, older adults displayed a more pronounced slowing in reaction times for correctly recalled medium-quality in contrast to high-quality memories (*Z* = 4.10, *p* <.001).

After indicating that a word was remembered during delayed recall, participants had to choose the second letter of the associated word out of four letter options to ensure correct word retrieval. Correct answers were thus possible by random selection of the correct letter option (25 % hits in case of random answers). To rule out that the comparably small effect of gaining an item was due to random guessing, we computed the probability of a correct letter selection after indicating that a word was remembered for all low-quality items. In both age groups this probability was significantly higher than 25 % (younger adults: *t*(29) = 9.86, *p* <.001, *M*_YA_ = 52.24 %; older adults: *t*(35)=5.75, *p* <.001, *M*_OA_ = 36.27 %). Hence, correct responses to low-quality items indicated an actual memory gain.

To sum up, we found comparable memory gains for both age groups, whereas memory maintenance was reduced in older adults. This reduction was most pronounced for memories of medium quality. Given that our analysis procedure was based on item-specific learning trajectories within single individuals, our procedure effectively controlled for inter-individual and age differences in memory encoding on Day 1. Hence, we suggest that the overnight changes in memory maintenance reflect age differences in sleep-dependent memory consolidation that vary with initial encoding strength.

### Sleep architecture in older adults shifts from SWS to lighter NREM sleep stages

Sleep architecture changes across the adult lifespan (Ohayon et al. 2004), but is to some extent also variable within individuals (Gais et al. 2002; Mölle et al. 2011) and can be shaped by learning experiences (Huber et al. 2004; Mölle et al. 2009). We thus first examined intra- as well as inter-individual variation in global sleep parameters. All sleep measures, except for sleep latency (i.e., the time needed to fall asleep) and the relative amount of REM sleep (both *p* ≥.053; both *F* ≤ 3.92), showed significant main effects of AGE (see Table 1). More precisely, we found the expected decreased proportion of SWS in older adults compensated for by an increase in the lighter NREM sleep stages 1, and 2. Neither the main effect of TIME nor the AGE-by-TIME interaction were significant for any of the variables, indicating that overall sleep architecture did not change as a result of the intense learning session on Day 1.

**Table 1.**
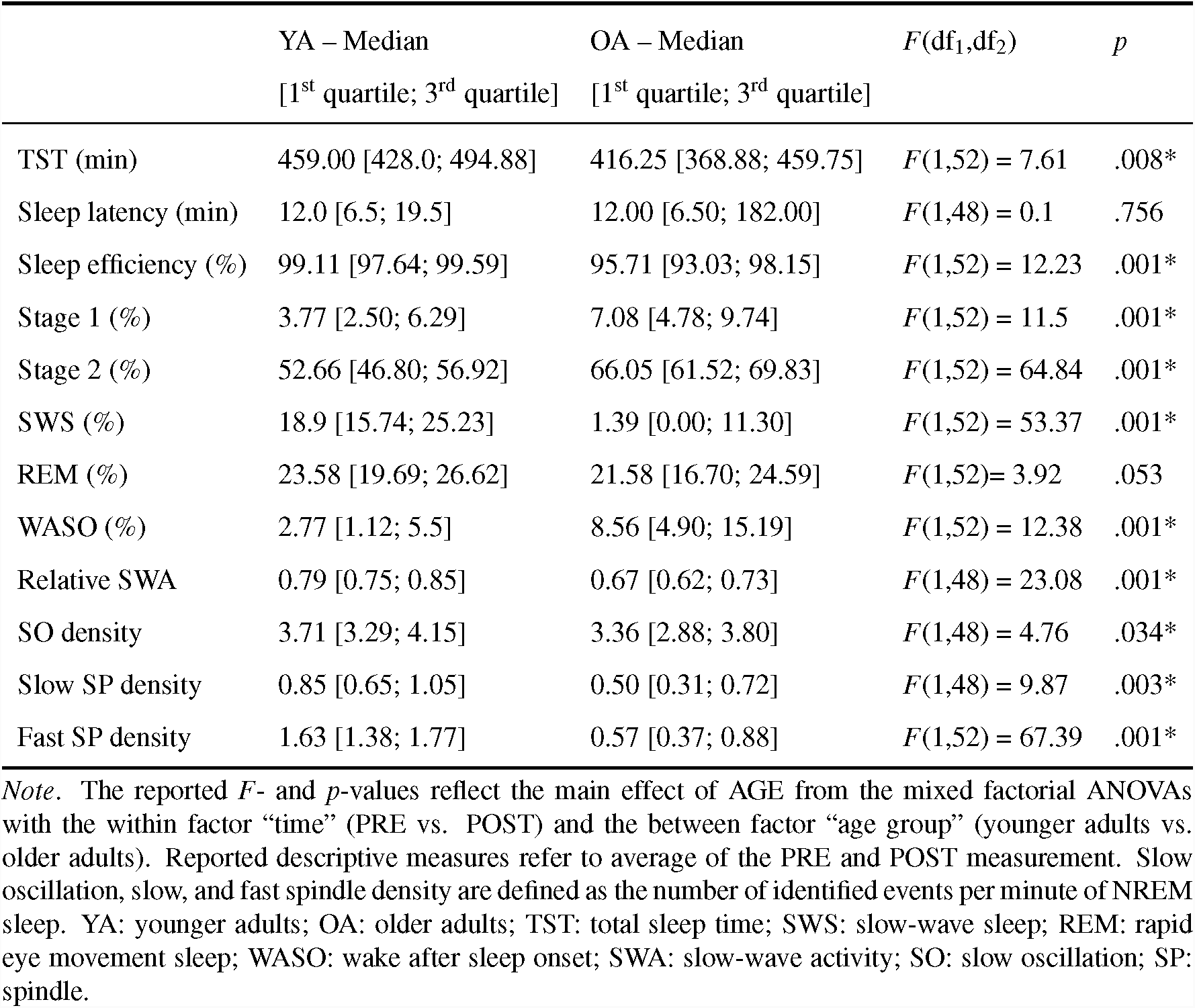
*Age-related changes in sleep variables*

Visual scoring of sleep stages is difficult in age-comparative studies. The fixed amplitude criteria for scoring SWS typically penalize older adults, as they generally show lower EEG amplitudes (Carrier et al. 2011). Accordingly, almost one third of our older sample failed to meet the amplitude criteria of SWS when visually scoring the PSG data (*n* = 9). Hence, we combined NREM sleep stages 2 and SWS to focus on more fine-grained physiological measures of NREM sleep. SWA, defined as relative power in the delta frequency range (0.5–4.5 Hz), slow oscillations (0.5–1 Hz), slow spindles (9–12.5 Hz), and fast spindles (12.5–16 Hz) were significantly reduced in older compared to younger adults (all *F* ≥ 4.76, all *p* ≤.034). SWA was the only variable showing a significant main effect of TIME (*F*(1,48) = 7.20, *p* <.001; *Md*_PRE_ = 0.7, *Md*_POST_ = 0.74). Intense learning distinctly boosted SWA. The overnight change in SWA, though, did not relate to any measure of memory consolidation (all |*r*| ≤.28, *p* ≥.051). As the AGE-by-TIME interaction was not significant for any other sleep measure, we restricted our analysis to sleep after learning only in the following.

### Senescent sleep does not drive inter-individual differences in memory consolidation

Aging causes broad alterations in sleep architecture and physiology (Mander et al. 2017). To account for the interdependency of sleep processes and their joint contributions to memory consolidation, we chose a multivariate approach to examine individual differences in aging as reflected in multiple physiological indicators of sleep. We then asked how these differences in aging relate to memory consolidation as captured by overnight memory maintenance and gain (Materials and Methods; see Supplementary Tables 1–3 for the bivariate sleep–memory associations). Using PLSC (Krishnan et al. 2011; Haenlein and Kaplan 2004), we identified a significant latent variable (LV) capturing reductions in frontal SWA, as well as in the density of frontal slow oscillations, frontal slow spindles, and central fast spindles with advancing age (*r*_AGE_ = 0.73, *p* <.001). Bootstrap ratios (BSR) indicated that all included sleep variables reliably contributed to the LV (all BSR ≤ –2.27; Figure 3a). The latent profile suggests that advancing age is accompanied by a simultaneous reduction in multiple neurophysiological markers of memory consolidation during NREM sleep. Given the nature of the extracted LV, older adults have a greater manifestation of the LV, that is, higher individual *latent sleep profile scores* (LSPS). The LSPS ranges considerably overlapped between younger and older adults (Figure 3b), suggesting that, although the individual sleep profile changes with aging, younger and older adults are not completely distinct in their sleep profiles.

**Figure 3.**
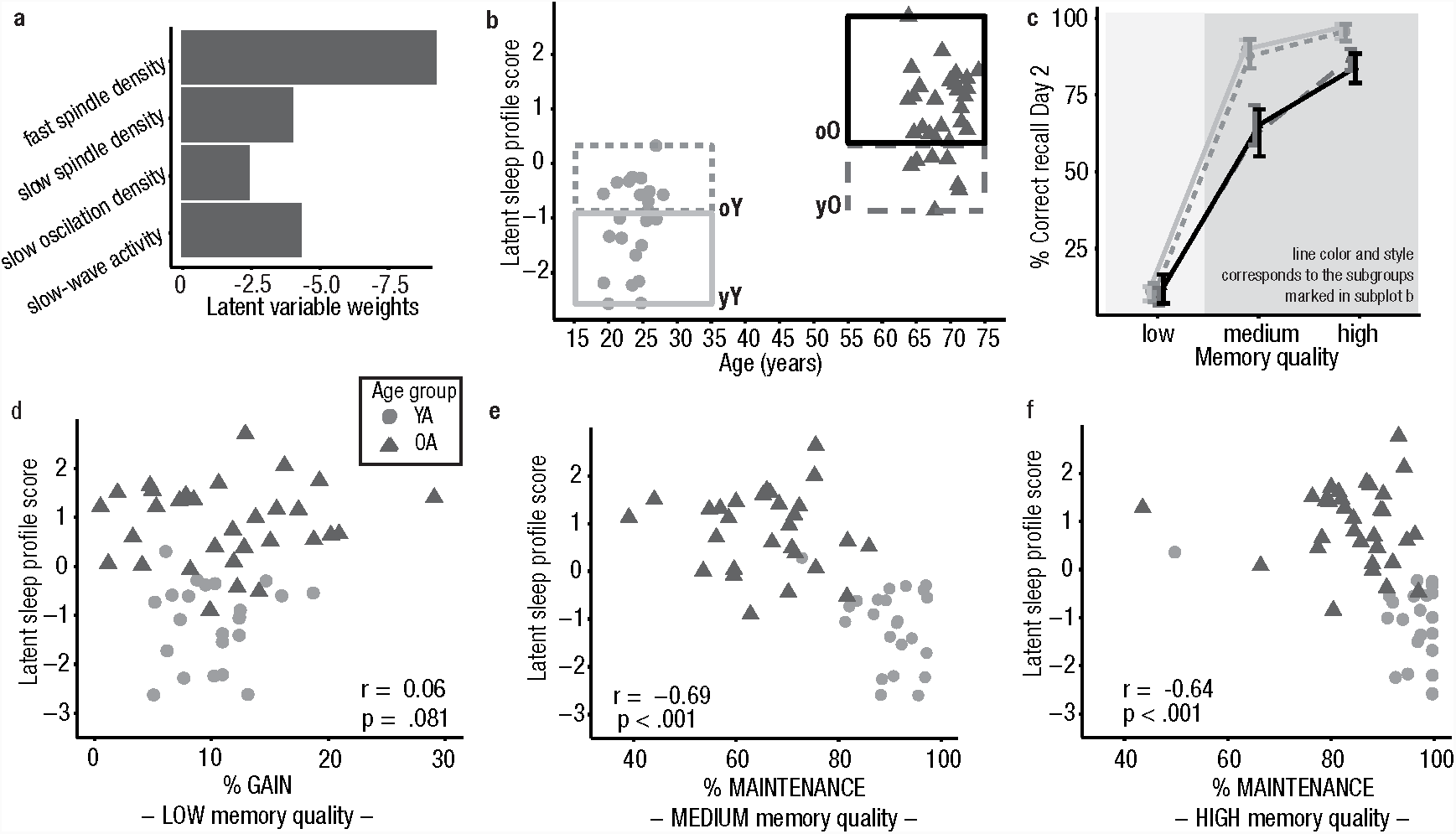
Sleep–memory associations in younger and older adults. (a) All sleep variables contribute to the latent variable capturing the common variance between participants’ age and sleep. Latent variable weights (in *Z*-scores) demonstrate that all sleep variables have a stable negative relation to age. (b) Each participant’s expression of the latent variable is plotted against age. Overlap between the age groups is marked by dashed gray boxes. Sleep in younger and older adults is not completely distinct. (c) Median behavioral performance for all subgroups (grouping and line style corresponding to b) is displayed. The first and third quartile is depicted as an error bar. Memory gain (shaded in light gray) is similar in all subgroups. Memory maintenance (shaded in darker gray) is modulated by the sleep profile but differs between younger and older adults, even when they have the same sleep profile. (d–f) Each participant’s latent sleep profile score is plotted against the behavioral measures. Spearman’s rank-order correlation coefficients for the whole sample are displayed. Maintenance of both medium- and high-quality memories relates to the latent sleep profile score across age groups. Note the ceiling performance in younger adults, in particular for high-quality memories. YA: younger adults, OA: older adults, yY: young–Young (= younger adults showing a clearly distinct profile from older adults), oY: old–Young (= younger adults exhibiting a sleep profile comparable to older adults), yO: young–Old (= older adults with a ‘youth-like’ sleep profile), oO: old–Old (=older adults with a sleep profile clearly distinct from younger adults).

Using Spearman’s rank-order correlation coefficient, we did not find a significant correlation between the LSPS and memory gain across both age groups (*r* = 0.06, *p* =.681, Figure 3d). In contrast, maintenance of medium- and high-quality memories was negatively related to the LSPS (*r*_medium_ = –.66, *p* <.001; *r*_high_ = –.64, *p* <.001, Figure 3e and f). The more participants displayed a senescent sleep profile (i.e., reduced SWA, slow oscillations, slow and fast spindles), the worse their memory maintenance was. When conducted separately within each age group, none of the correlations reached significance (all |*r*| ≤ 0.13, all *p* ≥.548, Supplementary Tables 2 and 3). Moreover, none of the single sleep measures was reliably associated with memory maintenance and gain in younger and older adults (all |*r*| ≤.41, all *p* ≥.049, α-level adjusted to .00028, Supplementary Tables 2 and 3).

If sleep, as previously suggested, is the major driving force behind age differences in memory maintenance, participants of different age groups with similar sleep profiles should be comparable in their ability to maintain memories between Day 1 and Day 2. Based on their LSPS, we identified those participants revealing a similar sleep profile (outlined boxes in Figure 3b). Based on this grouping we obtained 4 subgroups (Supplementary Table 4; young–Young [yY] = younger adults showing a clearly distinct sleep profile from older adults [*n* = 12]; old–Young [oY] = younger adults exhibiting a sleep profile comparable to older adults [*n* = 11]; young–Old [yO] = older adults with a ‘youth-like’ sleep profile [*n* = 7]; old–Old [oO] = older adults with a sleep profile clearly distinct from younger adults [*n* = 24]). Using a mixed factorial ANOVA with the between-subjects factor SUBGROUP and the within-subjects factor MEMORY QUALITY, we found significant main effects for both factors (MEMORY QUALITY: *F*(2, 100) = 1657.47, *p* <.001; SLEEP PROFILE SUBGROUP: *F*(3, 50) = 12.17, p <.001) along with a significant interaction (*F*(6, 100) = 18.75, *p* <.001; Figure 3c). Post-hoc tests indicated that within the two age groups, the classification according to sleep profile did not result in any significant behavioral differences (all |*Z*| ≤ 1.28, *p* ≥.202). Descriptively, memory maintenance of younger adults exhibiting a sleep profile similar to older adults (oY) was reduced compared to younger adults with a sleep profile clearly distinct from older adults (yY) (Figure 3c; Supplementary Table 4). For memories of medium quality, in spite of a comparable sleep profile, maintenance still differed significantly between younger and older adults (*Z* = 3.31, *p* <.001). For high-quality memories, though, this age effect disappeared when controlling for multiple testing (*Z* = 2.37, *p* =.018, α-level adjusted to.006).

To summarize, we found evidence that, in contrast to memory gain, age-related changes in sleep coincided with worse maintenance of medium- and high-quality memories. Memory maintenance might rely on processes during sleep that are impaired in aged individuals. However, we demonstrate that, within each age group, inter-individual differences in sleep patterns did not reliably account for inter-individual differences in memory consolidation. Moreover, the similarity of sleep patterns between younger and older adults did not reverse the effect of age on the consolidation of medium-quality memories. We thus conclude that a senescent sleep profile (characterized by reduced SWA, slow oscillations, slow, and fast spindles) does not account for all observed age-related reductions in memory maintenance.

### Age differences in brain structure coincide with sleep and memory impairments

Recently, a more comprehensive view on age-related changes in memory consolidation emerged by including measures of structural brain integrity (Varga et al. 2016; Mander et al. 2013; Helfrich et al. 2018; Muehlroth et al. in press). Again, we used PLSC to examine how age-related structural atrophy in various brain regions relates to the tendency to exhibit a senescent sleep profile and the ability to consolidate memories (cf. Materials and Methods; see Supplementary Tables 1–3 for the bivariate brain–memory associations). In analogy to the analysis described above, we used PLSC to extract a LV reflecting age-related reductions in gray matter volume in the mPFC, thalamus, entorhinal cortex, and the hippocampus (*r*_AGE_ = 0.78, *p* <.001). ROIs reliably contributed to the LV (all BSR ≤ –6.12; Figure 4a). The score quantifying each participant’s expression of the LV, further referred to as *latent brain structure score* (LBSS), was positively related to the LSPS (*r* = 0.65, *p* <.001; Supplementary Figure 2). Within each age group separately, however, this association was not significant (all |*r*| ≤.13, all *p* ≥.500; Supplementary Tables 2 and 3).

**Figure 4.**
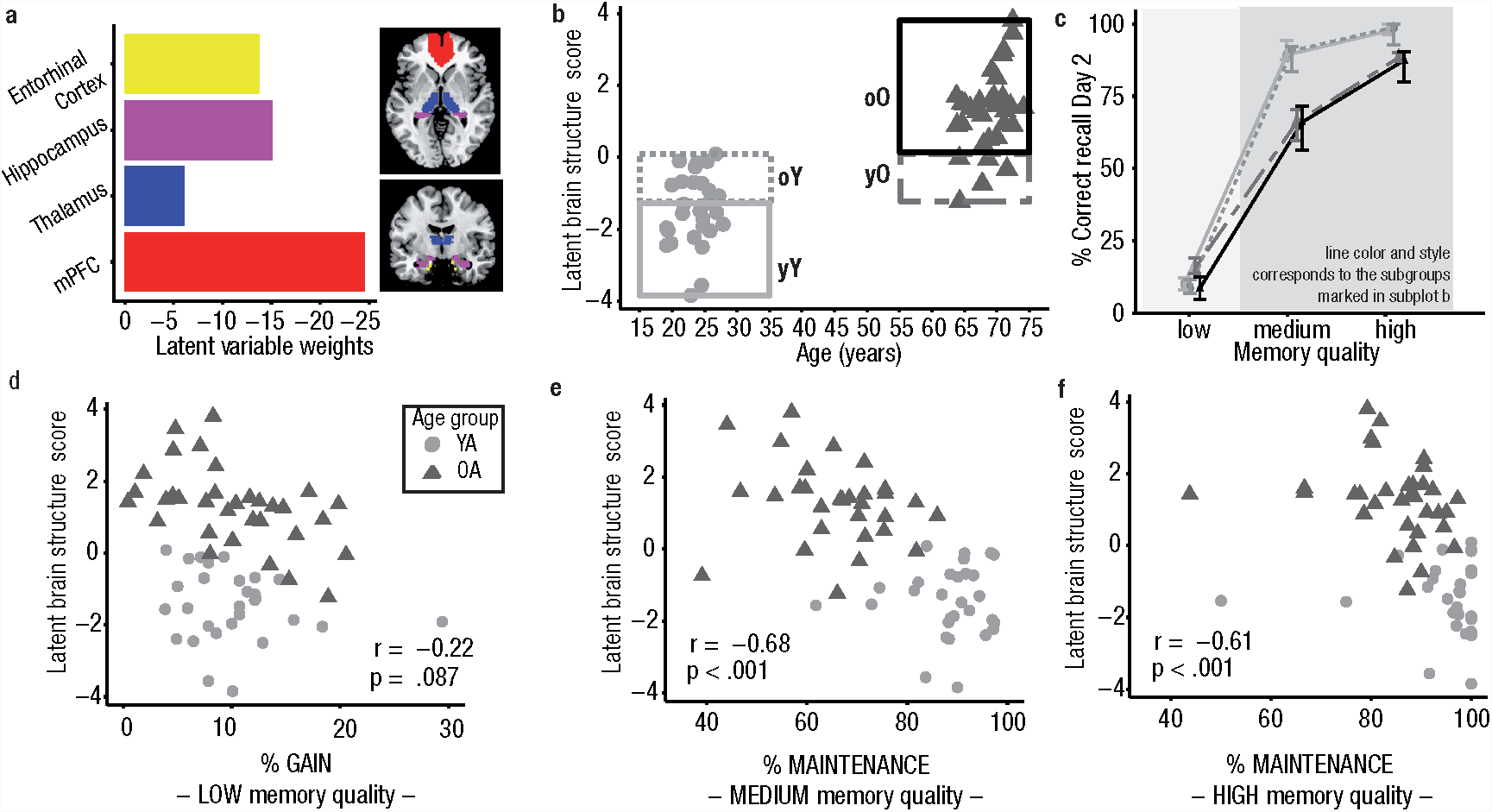
Brain–memory associations in younger and older adults. (a) Brain regions involved in slow wave and spindle generation and memory processing contribute to the latent variable capturing the common variance between participants’ age and brain structure (quantified using voxel-based morphometry). Latent variable weights (in *Z*-scores) demonstrate that all regions have a stable negative relation with age (all BSR ≤ –6.12). Bar colors correspond to the colors of the masked regions. (b) Each participant’s expression of the latent variable is plotted against age. Overlap between the age groups is indicated by dashed gray boxes. Younger and older participant groups with clearly differing brain structure are outlined by a solid line. Only few older adults show structural brain integrity comparable to younger adults. (c) Median behavioral performance for all subgroups is shown. The first and third quartile is depicted as an error bar. Memory gain (shaded in light gray) is similar in all subgroups. Memory maintenance (shaded in darker gray) differs between younger and older adults, even when they express the same structural brain integrity. Brain structure itself does not modulate behavior within age groups. (d–f) Each participant’s latent brain structure score plotted against the behavioral measures. Spearman’s rank-order correlation coefficients for the whole sample are displayed. Maintenance of both medium- and high-quality memories relates to the latent brain structure score across age groups. BSR: bootstrap ratio, YA: younger adults, OA: older adults, yY: young–Young (= younger adults showing a clearly distinct profile from older adults), oY: old–Young (= younger adults exhibiting a sleep profile comparable to older adults), yO: young–Old (= older adults with a ‘youth-like’ sleep profile), oO: old–Old (=older adults with a sleep profile clearly distinct from younger adults).

Maintenance of medium- and high-quality memories was negatively related to the LBSS (*r*_medium_ = –.68, *p* <.001; *r*_high_ = –.61, *p* <.001; Figure 4e and f). Nevertheless, these associations were only stable across age groups (within age groups all |*r*| ≤.31, all *p* ≥.08; Supplementary Tables 2 and 3). Although only on a trend level in the whole sample (*r* = –.22, *p* =.087), the correlation between the LBSS and memory gain reached significance in older, but not younger adults (*r*_OA_ = –.47, *p*_OA_ =.007; *r*_YA_ = –.2, *p*_YA_ =.302). Correlation coefficients, however, did not differ significantly between younger and older adults (*Z* = –1.13, *p* =.26).

As only a small proportion of older adults (*n* = 5) showed a structural brain integrity that was comparable to younger adults (Figure 4b), we could not follow the same rationale as described above to examine whether subjects in different age groups with similar structural brain integrity comparably maintained memories across sleep (but see Figure 4c and Supplementary Table 5, for an illustration of this effect). Compared to physiological markers of sleep, structural brain integrity separated the two age groups more distinctly (Figure 4b) but did not seem to differentiate behavior within age groups.

## Discussion

The present study asked whether variation in the encoding quality of individual associative memories determines the degree of age-related impairments in memory consolidation. We demonstrate that age differences in memory consolidation during sleep were maximized for the maintenance of mnemonic contents of medium encoding quality. By contrast, we did not find age differences in the proportion of associations gained overnight. It appears that age differences in sleep-dependent memory consolidation are confined to consolidation mechanisms that rely on intact NREM sleep. Using a multivariate approach, we successfully integrated patterns of sleep physiology and brain structure and, thus, overcame the typical use of multiple bivariate correlations. However, neither age differences in NREM sleep physiology nor in brain structure could fully account for the observed age-related reductions in overnight protection against forgetting.

### Encoding quality determines the extent of age differences in memory consolidation

Meeting the long demand to go beyond average net measures of memory (Tulving 1964, 1967), we separated different consolidation mechanisms based on the encoding quality of the respective memory (Dumay 2016; Fenn and Hambrick 2013). Our results are in line with ongoing discussions in the field, suggesting that the major role of sleep can be seen in the maintenance rather than the gain of memories (Dumay 2018; Fenn and Hambrick 2013; Nettersheim et al. 2015; Schreiner et al. 2018; but see Walker 2005).

By demonstrating that memory maintenance is differentially affected by aging, with most age-related deficits for memories of medium encoding quality, we underpin the notion of an active consolidation process that stabilizes mnemonic contents selectively (Diekelmann et al. 2009; Drosopoulos et al. 2007; Kuriyama et al. 2004; Schapiro et al. 2018; Stickgold 2010; Stickgold and Walker 2013; Tucker and Fishbein 2008). Already establishing strong robust memory representations during encoding might render subsequent consolidation processes redundant (Schoch et al. 2017). Memories of intermediate quality, in contrast, might have the necessary but not yet sufficient strength that prioritizes them for subsequent active consolidation mechanisms that include active processes like the reactivation and redistribution of memory traces (Schapiro et al. 2018; Stickgold 2010). Since these processes are particularly disrupted in old age (Cordi et al. 2018; Gerrard et al. 2008; Helfrich et al. 2018), pronounced age-related deficits in memory, as observed here, might be the consequence of age-related impairments in the reactivation and redistribution of memory traces.

Memory gain, in contrast, was comparable across age groups. The fact that memories that could not be recalled on Day 1 were readily available on Day 2 may point to an active consolidation mechanism that raises memory representations above a pre-sleep learning threshold (Walker 2005). However, although successful recall of a memory indicates the existence of a reliable memory representation, unsuccessful recall does not necessarily indicate the opposite. Reduced attentional resources (Anderson et al. 2000; Craik et al. 2010) and incomplete retrieval search (Grady and Craik 2000; Raaijmakers and Shiffrin 1981), which are both known to affect retrieval in older adults, might have resulted in failed pre-sleep memory retrieval on Day 1 despite the availability of the respective memory trace (Habib and Nyberg 2007). Moreover, retrieval itself can result in memory suppression, a phenomenon called “retrieval-induced forgetting” (Bäuml and Kliegl 2017; MacLeod and Macrae 2001). Importantly, retrieval-induced forgetting is temporary and recovers over an interval of 24 hours (Abel and Bäuml 2014; MacLeod and Macrae 2001). This is equivalent to the interval used in this study (but see Abel and Bäuml 2012). In line with the similar memory gain in younger and older adults observed here, retrieval-induced forgetting has also been shown to be independent of aging (Hogge et al. 2008). We thus speculate that the effect of memory gain in our analyses reflects the inaccessibility of specific memories in the final recall on Day 1 (despite general availability of the respective trace) rather than a memory improvement across sleep.

To conclude, our findings demonstrate that by taking into account unknown variation in the quality of individual memories, differential effects of aging on memory consolidation can be explained. When studying age differences in sleep-dependent memory consolidation, a tight control of learning success is thus demanded.

### Sleep physiology and brain structure alone do not account for age differences in memory maintenance

Prominent age-related changes in sleep physiology that particularly affect NREM sleep (Carrier et al. 2011; Mander et al. 2017; Ohayon et al. 2004), are assumed to constitute one of the key mechanisms causing consolidation deficits in old age. Indeed, older adults in our sample exhibited reductions in SWA as well as slow oscillations, slow, and fast spindle density. Moreover, age-related reductions in the expression of rhythmic neural activity during NREM sleep in older adults coincided with worse maintenance of medium- and high-quality memories (but not overnight memory gain). The reduced presence of slow waves, slow oscillations, and spindles may thus indicate impairments of the coordinated dialogue between the hippocampus and neocortex that hinder the transfer of labile memory representations to permanent stores in the neocortex (Helfrich et al. 2018; Mander et al. 2013; Muehlroth et al. in press; Varga et al. 2016). However, equating younger and older adults with regard to their sleep profiles did not entirely remove age differences in memory maintenance. Also within age groups, inter-individual differences in the extent of the expression of an ‘aged’ sleep profile did not account for inter-individual differences in either memory gain or maintenance. Hence, we suggest that inter-individual differences in NREM sleep physiology may not qualify as the sole source of age differences in memory maintenance observed in the present study.

Age-related changes in sleep physiology may be driven by senescent changes in the structural integrity of brain areas involved in slow wave and spindle generation (Saletin et al. 2013). Indeed, previous reports suggested age-related gray matter atrophy as the main cause of changes in sleep with advancing age (Dubé et al. 2015; Fogel et al. 2017; Landolt and Borbély 2001; Mander et al. 2013; Varga et al. 2016). Although most pronounced in frontal brain areas (Giorgio et al. 2010), aging also results in loss of gray matter volume in memory-relevant regions in the MTL (Raz et al. 2005). In combination, these wide-spread structural alterations in sleep and memory networks might impact the hippocampal reactivation of memory traces during sleep and their redistribution among brain systems (Cordi et al. 2018; Gerrard et al. 2008; Helfrich et al. 2018; Mander et al. 2013; Muehlroth et al. in press).

In general, we observed that participants with reduced structural brain integrity exhibit a bmore ‘aged’ sleep profile and worse maintenance of both medium- and high-quality memories. However, correlations between brain structure, sleep, and memory only held across, but not within age groups. Similar to our analysis of age-related differences in sleep physiology and their relation to overnight memory maintenance, we do not find strong support for the assumption that changes in structural brain integrity are the major predictor for inter-individual variation in sleep and memory consolidation.

### Towards a mechanistic understanding of age-related alterations in memory consolidation during sleep

The present study provides cross-sectional evidence that age differences in NREM sleep physiology and structural brain integrity contribute to age differences in sleep-dependent memory consolidation (Mander et al. 2013; Varga et al. 2016). However, our results raise doubts that inter-individual differences in sleep physiology can sufficiently predict the success of memory consolidation during sleep. Thus, with the currently used methodology, it might be too early to consider sleep a possible biomarker of age-related pathological and non-pathological memory decline (Mander et al. 2016).

The importance of slow oscillations and spindles for memory consolidation has been underscored by a variety of studies experimentally manipulating sleep physiology (Marshall et al. 2006, 2004; Ngo et al. 2013; Van Der Werf et al. 2009) and by studies showing the benefit of targeted memory reactivation during periods of NREM sleep (Rasch et al. 2007). Nevertheless, correlational studies sometimes fail to detect similar relationships (Ackermann et al. 2015; Rasch and Born 2013, see Mantua 2018 for a recent discussion). Crucially, this does not contradict the important role of NREM sleep physiology for memory consolidation. Rather, it points to the fact that inter-individual differences in the mere *occurrence* of slow waves and sleep spindles might be an insufficient predictor of between-person differences in memory consolidation. This view is supported by recent reports suggesting that consolidation success relies on the fine-tuned *coordination* of rhythmic neural events during NREM sleep rather than their simple presence (Latchoumane et al. 2017; Maingret et al. 2016). Along these lines, dispersed slow oscillation–spindle coupling could predict consolidation impairments in older adults (Helfrich et al. 2018; Muehlroth et al. in press). Hence, we suggest that a mechanistic understanding of the causes and consequences of age differences in memory consolidation ultimately requires novel analytic tools that disclose the fine-tuned interplay between rhythmic neural events during NREM sleep, the interaction of different brain structures or the interplay of brain structure, and sleep oscillations. This will finally pave the road for novel therapeutic interventions that can reveal their full therapeutic capability to reduce or delay cognitive decline in old age.

## Acknowledgements

This study was conducted within the *‘Cognitive and Neural Dynamics of Memory across the Lifespan (ConMem)’* project at the Center for Lifespan Psychology, Max Planck Institute for Human Development. We thank Maren J. Cordi for helping us to set up the technical equipment, Xenia Grande for organizing data collection, Kristina Günther for help in participant recruitment, Julia Delius for editorial assistance, and all student assistants of the *ConMem* project collecting the data. We are grateful to all members of the *ConMem* project for helpful feedback on the analysis. Finally, we thank all study participants for their time.

