## Supplemental Information for "Memory quality modulates the effect of aging on memory consolidation during sleep: Reduced maintenance but intact gain"

---

*Supplementary Table 1. Bivariate correlations between sleep variables, brain regions of interest, and memory measures in the* ***whole sample***

| | | *1* | *2* | *3* | *4* | *5* | *6* | *7* | *8* | *9* | *10* | *11* | *12* | *13* |
|---|---|---|---|---|---|---|---|---|---|---|---|---|---|---|
| *1* | *Fast SP density* | | | | | | | | | | | | | |
| *2* | *Slow SP density* | **0.517** ***(<.001)*** | | | | | | | | | | | | |
| *3* | *SO density* | 0.183 *(.182)* | –0.132 *(.335)* | | | | | | | | | | | |
| *4* | *SWA* | 0.285 *(.035)* | 0.059 *(.669)* | **0.458** ***(<.001)*** | | | | | | | | | | |
| *5* | *mPFC* | **0.583** ***(<.001)*** | **0.434** ***(<.001)*** | 0.229 *(.098)* | **0.481** ***(<.001)*** | | | | | | | | | |
| *6* | *Thalamus* | 0.440 *(.001)* | 0.390 *(.004)* | –0.065 *(.644)* | 0.160 *(.253)* | **0.635** ***(<.001)*** | | | | | | | | |
| *7* | *Hippocampus* | **0.556** ***(<.001)*** | 0.313 *(.023)* | 0.074 *(.595)* | 0.413 *(.002)* | **0.726** ***(<.001)*** | **0.662** ***(<.001)*** | | | | | | | |
| *8* | *Entorhinal cortex* | **0.489** ***(<.001)*** | 0.372 *(.006)* | 0.052 *(.712)* | 0.393 *(.004)* | **0.616** ***(<.001)*** | **0.548** ***(<.001)*** | **0.800** ***(<.001)*** | | | | | | |
| *9* | *Age* | **–0.682** ***(<.001)*** | **–0.460** ***(<.001)*** | –0.180 *(.194)* | **–0.426** ***(<.001)*** | **–0.758** ***(<.001)*** | **–0.603** ***(<.001)*** | **–0.722** ***(<.001)*** | **–0.666** ***(<.001)*** | | | | | |
| *10* | *Low quality (gain)* | –0.013 *(.925)* | 0.028 *(.839)* | –0.071 *(.607)* | –0.160 *(.243)* | 0.108 *(.400)* | 0.318 *(.011)* | 0.142 *(.267)* | 0.131 *(.306)* | –0.088 *(.488)* | | | | |
| *11* | *Medium quality (maintenance)* | **0.674** ***(<.001)*** | 0.337 *(.012)* | 0.274 *(.043)* | 0.396 *(.003)* | **0.665** ***(<.001)*** | **0.568** ***(<.001)*** | **0.584** ***(<.001)*** | **0.527** ***(<.001)*** | **–0.766** ***(<.001)*** | 0.264 *(.032)* | | | |
| *12* | *High quality (maintenance)* | **0.512** ***(<.001)*** | 0.338 *(.012)* | 0.329 *(.014)* | 0.360 *(.007)* | **0.610** ***(<.001)*** | **0.575** ***(<.001)*** | **0.458** ***(<.001)*** | **0.406** ***(<.001)*** | **–0.688** ***(<.001)*** | 0.325 *(.008)* | **0.808** ***(<.001)*** | | |
| *13* | *LSPS* | **–0.867** ***(<.001)*** | **–0.572** ***(<.001)*** | –0.419 *(.002)* | **–0.610** ***(<.001)*** | **–0.712** ***(<.001)*** | **–0.414** ***(<.001)*** | **–0.575** ***(<.001)*** | **–0.533** ***(<.001)*** | **0.727** ***(<.001)*** | 0.057 *(.680)* | **–0.689** ***(<.001)*** | **–0.640** ***(<.001)*** | |
| *14* | *LBSS* | **–0.609** ***(<.001)*** | –0.429 *(.002)* | –0.090 *(.525)* | –0.421 *(.002)* | **–0.870** ***(<.001)*** | **–0.813** ***(<.001)*** | **–0.919** ***(<.001)*** | **–0.848** ***(<.001)*** | **0.775** ***(<.001)*** | –0.219 *(.087)* | **–0.680** ***(<.001)*** | **–0.608** ***(<.001)*** | **0.653** ***(<.001)*** |

*Note*. Spearman's correlation coefficients. Correlations in black fall below an α-level of .05. Bold black correlation coefficients fall below the Bonferroni-adjusted α-level of .00028. LBSS: latent brain structure score, LSPS: latent sleep profile score, mPFC: medial prefrontal cortex, SP: spindle, SO: slow oscillation, SWA: slow-wave activity.

*Supplementary Table 2. Bivariate correlations between sleep variables, brain regions of interest, and memory measures in* ***younger adults***

| | | *1* | *2* | *3* | *4* | *5* | *6* | *7* | *8* | *9* | *10* | *11* | *12* | *13* |
|---|---|---|---|---|---|---|---|---|---|---|---|---|---|---|
| *1* | *Fast SP density* | | | | | | | | | | | | | |
| *2* | *Slow SP density* | 0.661 *(.001)* | | | | | | | | | | | | |
| *3* | *SO density* | –0.447 *(.030)* | –0.107 *(.618)* | | | | | | | | | | | |
| *4* | *SWA* | –0.333 *(.112)* | –0.275 *(.193)* | 0.503 *(.013)* | | | | | | | | | | |
| *5* | *mPFC* | 0.042 *(.835)* | 0.064 *(.765)* | –0.066 *(.759)* | –0.013 *(.953)* | | | | | | | | | |
| *6* | *Thalamus* | 0.349 *(.075)* | 0.250 *(.237)* | –0.351 *(.093)* | –0.379 *(.069)* | 0.444 *(.015)* | | | | | | | | |
| *7* | *Hippocampus* | 0.093 *(.642)* | –0.210 *(.324)* | –0.366 *(.079)* | –0.283 *(.180)* | 0.322 *(.083)* | 0.477 *(.008)* | | | | | | | |
| *8* | *Entorhinal cortex* | 0.209 *(.293)* | 0.154 *(.471)* | –0.237 *(.265)* | –0.138 *(.518)* | 0.022 *(.910)* | 0.342 *(.065)* | 0.483 *(.007)* | | | | | | |
| *9* | *Age* | –0.441 *(.025)* | –0.370 *(.083)* | 0.308 *(.152)* | 0.023 *(.919)* | –0.081 *(.676)* | –0.129 *(.503)* | –0.067 *(.731)* | –0.111 *(.564)* | | | | | |
| *10* | *Low quality (gain)* | –0.008 *(.967)* | 0.306 *(.146)* | –0.134 *(.531)* | –0.407 *(.049)* | 0.176 *(.351)* | 0.232 *(.217)* | 0.060 *(.752)* | 0.155 *(.414)* | 0.153 *(.427)* | | | | |
| *11* | *Medium quality (maintenance)* | 0.207 *(.299)* | 0.365 *(.080)* | 0.086 *(.688)* | –0.163 *(.444)* | –0.017 *(.929)* | 0.200 *(.287)* | –0.244 *(.193)* | –0.190 *(.313)* | –0.359 *(.057)* | 0.305 *(.101)* | | | |
| *12* | *High quality (maintenance)* | 0.127 *(.527)* | 0.358 *(.086)* | 0.096 *(.657)* | 0.002 *(.993)* | 0.179 *(.344)* | 0.279 *(.136)* | –0.248 *(.187)* | –0.132 *(.486)* | –0.382 *(.041)* | 0.219 *(.244)* | 0.535 *(.002)* | | |
| *13* | *LSPS* | –0.593 *(.003)* | –0.588 *(.041)* | –0.218 *(.315)* | –0.396 *(.062)* | –0.065 *(.767)* | 0.054 *(.806)* | 0.322 *(.134)* | –0.016 *(.944)* | 0.259 *(.232)* | 0.094 *(.670)* | –0.114 *(.604)* | –0.132 *(.548)* | |
| *14* | *LBSS* | –0.247 *(.223)* | –0.079 *(.719)* | 0.328 *(.127)* | 0.239 *(.271)* | **–0.620** ***(<.001)*** | **–0.808** ***(<.001)*** | **–0.841** ***(<.001)*** | **–0.628** ***(<.001)*** | 0.178 *(.353)* | –0.198 *(.302)* | 0.052 *(.790)* | –0.049 *(.802)* | –0.079 *(.719)* |

*Note*. Spearman's correlation coefficients. Correlations in black fall below an α-level of .05. Bold black correlation coefficients fall below the Bonferroni-adjusted α-level of .00028. LBSS: latent brain structure score, LSPS: latent sleep profile score, mPFC: medial prefrontal cortex, SP: spindle, SO: slow oscillation, SWA: slow-wave activity.

*Supplementary Table 3. Bivariate correlations between sleep variables, brain regions of interest, and memory measures in* ***older adults***

| | | *1* | *2* | *3* | *4* | *5* | *6* | *7* | *8* | *9* | *10* | *11* | *12* | *13* |
|---|---|---|---|---|---|---|---|---|---|---|---|---|---|---|
| *1* | *Fast SP density* | | | | | | | | | | | | | |
| *2* | *Slow SP density* | 0.306<br>*(.094)* | | | | | | | | | | | | |
| *3* | *SO density* | 0.151<br>*(.415)* | –0.402<br>*(.026)* | | | | | | | | | | | |
| *4* | *SWA* | –0.114<br>*(.541)* | –0.107<br>*(.566)* | 0.312<br>*(.087)* | | | | | | | | | | |
| *5* | *mPFC* | –0.052<br>*(.788)* | 0.232<br>*(.225)* | 0.127<br>*(.511)* | 0.033<br>*(.867)* | | | | | | | | | |
| *6* | *Thalamus* | –0.006<br>*(.977)* | 0.133<br>*(.488)* | –0.142<br>*(.461)* | –0.039<br>*(.839)* | 0.364<br>*(.038)* | | | | | | | | |
| *7* | *Hippocampus* | 0.101<br>*(.599)* | 0.160<br>*(.405)* | –0.027<br>*(.889)* | 0.065<br>*(.739)* | 0.228<br>*(.201)* | 0.457<br>*(.008)* | | | | | | | |
| *8* | *Entorhinal cortex* | –0.068<br>*(.723)* | 0.029<br>*(.881)* | –0.196<br>*(.308)* | 0.023<br>*(.905)* | 0.082<br>*(.648)* | 0.183<br>*(.306)* | **0.648**<br>***(<.001)*** | | | | | | |
| *9* | *Age* | –0.143<br>*(.444)* | –0.130<br>*(.486)* | 0.060<br>*(.747)* | 0.131<br>*(.482)* | –0.171<br>*(.341)* | –0.374<br>*(.032)* | –0.380<br>*(.029)* | –0.198<br>*(.270)* | | | | | |
| *10* | *Low quality (gain)* | 0.052<br>*(.779)* | –0.063<br>*(.734)* | –0.049<br>*(.794)* | –0.005<br>*(.978)* | 0.252<br>*(.156)* | 0.489<br>*(.004)* | 0.324<br>*(.067)* | 0.194<br>*(.278)* | –0.409<br>*(.013)* | | | | |
| *11* | *Medium quality (maintenance)* | 0.338<br>*(.063)* | –0.281<br>*(.126)* | 0.097<br>*(.603)* | –0.128<br>*(.492)* | –0.021<br>*(.908)* | 0.338<br>*(.054)* | 0.234<br>*(.189)* | 0.030<br>*(.867)* | –0.255<br>*(.133)* | 0.463<br>*(.004)* | | | |
| *12* | *High quality (maintenance)* | 0.118<br>*(.527)* | –0.169<br>*(.362)* | 0.218<br>*(.237)* | –0.017<br>*(.928)* | 0.202<br>*(.259)* | 0.469<br>*(.006)* | 0.228<br>*(.201)* | 0.003<br>*(.987)* | –0.358<br>*(.032)* | 0.502<br>*(.002)* | **0.620**<br>***(<.001)*** | | |
| *13* | *LSPS* | **–0.690**<br>***(<.001)*** | –0.426<br>*(.018)* | –0.379<br>*(.036)* | –0.408<br>*(.023)* | –0.203<br>*(.290)* | –0.012<br>*(.950)* | –0.237<br>*(.214)* | 0.090<br>*(.641)* | 0.114<br>*(.541)* | 0.008<br>*(.968)* | –0.004<br>*(.982)* | –0.094<br>*(.612)* | |
| *14* | *LBSS* | –0.033<br>*(.867)* | –0.180<br>*(.348)* | 0.073<br>*(.706)* | –0.049<br>*(.799)* | –0.554<br>*(.001)* | **–0.640**<br>***(<.001)*** | **–0.789**<br>***(<.001)*** | **–0.694**<br>***(<.001)*** | 0.278<br>*(.117)* | –0.465<br>*(.007)* | –0.260<br>*(.144)* | –0.311<br>*(.078)* | -0.130<br>*(.500)* |

*Note*. Spearman's correlation coefficients. Correlations in black fall below an $\alpha$-level of .05. Bold black correlation coefficients fall below the Bonferroni-adjusted $\alpha$-level of .00028. LBSS: latent brain structure score, LSPS: latent sleep profile score, mPFC: medial prefrontal cortex, SP: spindle, SO: slow oscillation, SWA: slow-wave activity.

*Supplementary Table 4. Memory gain and maintenance by sleep profile subgroup*

| | SLEEP PROFILE | | | |
|---|---|---|---|---|
| | Younger adults | | Older adults | |
| | young–Young ($n$ = 12) | old–Young ($n$ = 11) | young–Old ($n$ = 7) | old–Old ($n$ = 24) |
| GAIN | | | | |
| low-quality memory | 10.11 [7.11; 11.13] | 8.76 [6.91; 12.53] | 9.64 [4.05; 11.26] | 11.36 [6.09; 15.71] |
| MAINTENANCE | | | | |
| medium-quality memory | 91.55 [89.73; 94.71] | 89.29 [85.29; 94.95] | 62.82 [59.57; 72.92] | 66.35 [55.8; 71.47] |
| high-quality memory | 98.88 [95.00; 100.00] | 97.62 [94.39; 100.00] | 88.43 [84.57; 91.72] | 85.48 [80.27; 90.1] |

*Note*. Sleep profile subgroups correspond to the subgroups marked in Figure 3b that pinpoint younger and older adults with comparable and distinct sleep profiles: young–Young (= younger adults showing a clearly distinct sleep profile from older adults), old–Young (= younger adults exhibiting a sleep profile comparable to older adults), young–Old (= older adults with a 'youth-like' sleep profile), old–Old (=older adults with a sleep profile clearly distinct from younger adults).

*Supplementary Table 5. Memory gain and maintenance by brain structure subgroup*

| | BRAIN STRUCTURE | | | |
|---|---|---|---|---|
| | Younger adults | | Older adults | |
| | young–Young ($n$ = 17) | old–Young ($n$ = 12) | young–Old ($n$ = 5) | old–Old ($n$ = 28) |
| GAIN | | | | |
| low-quality memory | 9.60 [7.86; 11.72] | 8.57 [5.86; 10.3] | 15.28 [13.51; 18.9] | 8 [4.64; 11.91] |
| MAINTENANCE | | | | |
| medium-quality memory | 89.29 [87.76; 94.40] | 90.71 [83.45; 92.05] | 66.04 [59.52; 70.42] | 65.97 [56.36; 71.47] |
| high-quality memory | 97.62 [96.42; 100.00] | 98.91 [92.72; 100.00] | 88.43 [87.23; 90.00] | 87.41 [79.81; 90.5] |

*Note*. Brain structure subgroups correspond to the subgroups marked in Figure 4b that pinpoint younger and older adults with comparable and distinct brain structure profiles: young–Young (= younger adults with structural brain integrity clearly distinct from older adults), old–Young (= younger adults exhibiting a brain structure profile comparable to older adults), young–Old (= older adults with ‘youth-like’ structural brain integrity), old–Old (=older adults with a brain structure profile clearly distinct from younger adults).

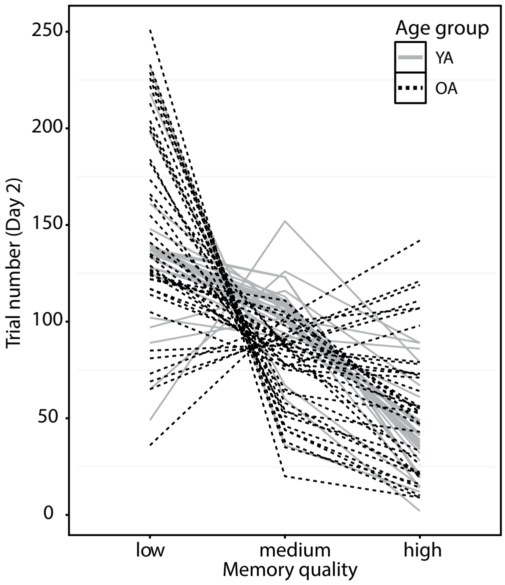


*Supplementary Figure 1*. Trial composition on Day 2. The number of trials during delayed retrieval on Day 2 is displayed for each memory quality condition (as defined by recall success on Day 1). Lines represent single subjects. Note the great inter-individual variability in trial composition on Day 2, which is caused by differential learning trajectories on Day 1. YA: younger adults, OA: older adults.

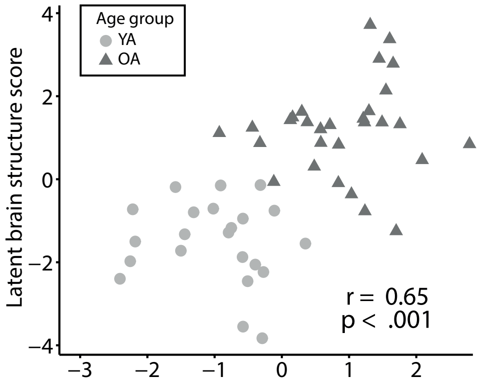


*Supplementary Figure 2*. Latent variable association. Each participant's latent brain structure score is plotted against the latent sleep profile score. *Spearman's* rank-order correlation coefficient for the whole sample is displayed. YA: younger adults, OA: older adults.
